# Circadian rhythm of temporal reproduction as a function of stimulus eccentricity in the visual field

**DOI:** 10.1101/2022.03.06.483173

**Authors:** Wei Wang, Xuanzi Yin, Yan Bao

## Abstract

The aim of this study was to elucidate how temporal reproduction changed in a diurnal time course. The overestimated reproduction showed a smaller difference to the standard in the morning and evening and a larger difference in the afternoon, and the underestimated reproduction showed an inverse change. No eccentricity effect in temporal production was observed, which indicates homogenous temporal processing shared in the perifoveal and peripheral visual fields.

---

Duration processing has been discussed on different time scales from the integration window of 100 -150 ms, 1 s to 2-3 s. The importance of circadian system has been identified in many cognitive domains. Various performance rhythms are possibly driven by different circadian oscillators, observable with distinct phase relationships (Lo, Groeger et al. 2012) under defined entrainment conditions. The circadian rhythm of a duration-specific process is still unclear, in particular temporal reproduction. Studies have shown that targets presented at the perifoveal visual field are more efficiently than at peripheral visual field (Carrasco, Evert et al. 1995, Wolfe, O’Neill et al. 1998). The eccentricity effect shows a stronger inhibition of return magnitude at the peripheral visual field compared to the perifoveal visual field (Bao, Lei et al. 2013). However, the eccentricity effect of temporal reproduction is rarely studied. In this study, we aimed to elucidate how temporal reproduction with different stimulus eccentricities changed in a diurnal time course. Based on the symmetry of the anatomical structure of visual pathway, only the left monocular visual field was employed in our study. The research was approved by the ethical committee of the University of Munich and was conducted in accordance with the principles of the Declaration of Helsinki. Written informed consent was obtained from all participants. Eighteen young healthy female participants of intermediate chronotype (MSFsc=04:21± 00:10; MEQ= 51± 0.99) were matched according to age (27.17± 0.49 years), BMI (20.24 ± 0.45 kg/m^2^), educational level (20.31± 2.09 years), depression-anxiety level (SDS: 44.22 ± 0.87; SAS: 35.39 ± 0.70) and sleeping time (workday: 23:23 ± 00:08 to 07:34 ± 00:08, freeday: 23:53 ± 00:08 to 08:46 ± 00:13). Participants were required to live strictly sleep-wake schedule for 1 week before the study. Afterwards, they reported to the isolated sleep room for two consecutive nights. The wake/sleep cycle is 17h/7h with the lighting illumination of 1000 lux/ 0 lux. The laboratory protocol started at 7:00 after the first adaptation night. A modified visual reproduction paradigm was adopted. Two standard durations (1500 ms and 4500 ms) randomly presenting at two stimulus eccentricities (7° and 21°) were asked to be reproduced during the waking time with 2-hour bins (7:00, 9:00, 11:00, 13:00, 15:00, 17:00, 19:00, 21:00, 23:00). Only left visual field and left eye was stimulated. Each trial started with a duration of 1000 ms of a white fixation cross (1°) centered on the black screen. Participants were required to wear an eye pad to mask their right eye, and seated approximately 48 cm in front of the monitor. They were instructed that they would see a white frame lasting for a certain amount of time. Afterwards they would see a white square lasting until they pressed a button on the computer keyboard. They were told to press the button when they thought the same amount of time had elapsed as for the standard stimulus. Participants were instructed not to count and were instructed to maintain their fixation at the central cross without shifting their gaze during each trial. An intertrial interval of 1000 ms with a blank screen was presented after the participant responds. Each condition was presented 15 times, leading to a total of 60 trials. The total task was divided into 6 sessions. A short break of 20 s between each session was inserted.

A 2 (duration) by 2 (stimulus eccentricity) by 9 (clock) general linear model repeated ANOVA for reproduced durations revealed no significant three-way interaction (F (8, 136) =1.757, *p* =0.091, 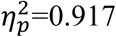) and none of the two-way interactions was significant (all *p* >0.05). The results showed significant main effects of duration and clock, F (1, 17) = 187.162, *p* <0.001, 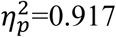, F (8, 136) = 3.398, *p* =0.001, 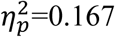, respectively. The absolute reproduced differences to the standards had a significant negative correlation, *r* = -0.6758, *p*=0.0457. Neither error ellipse overlaped the pole analyzing by Cosinor (Heart 2019), both reproduction thus were characterized by significant circadian rhythms (*p* <0.001, Figure 1. B). The Cosinor analysis showed that the MESOR of the reproduction of 4500ms (M=3750.76) was higher than of 1500 ms (M=2005.10). As to the zero-amplitude-test, the reproduction of 1500 ms (A=197.78, 95% CI was from 118.75 to 278.39) had a higher amplitude than of 4500 ms (A=189.12, 95% CI was from 49.10 to 329.33). Although the reproduction of 4500 ms (clock time of acrophase was 15:34) showed approximately 1 hour of acrophase ahead to 1500 ms (clock time of acrophase was 16:32). The overlap of 95% confidence arcs indicated no statistically significant difference was found between the circadian acrophases in the two reproductions.

**Figure 1.**
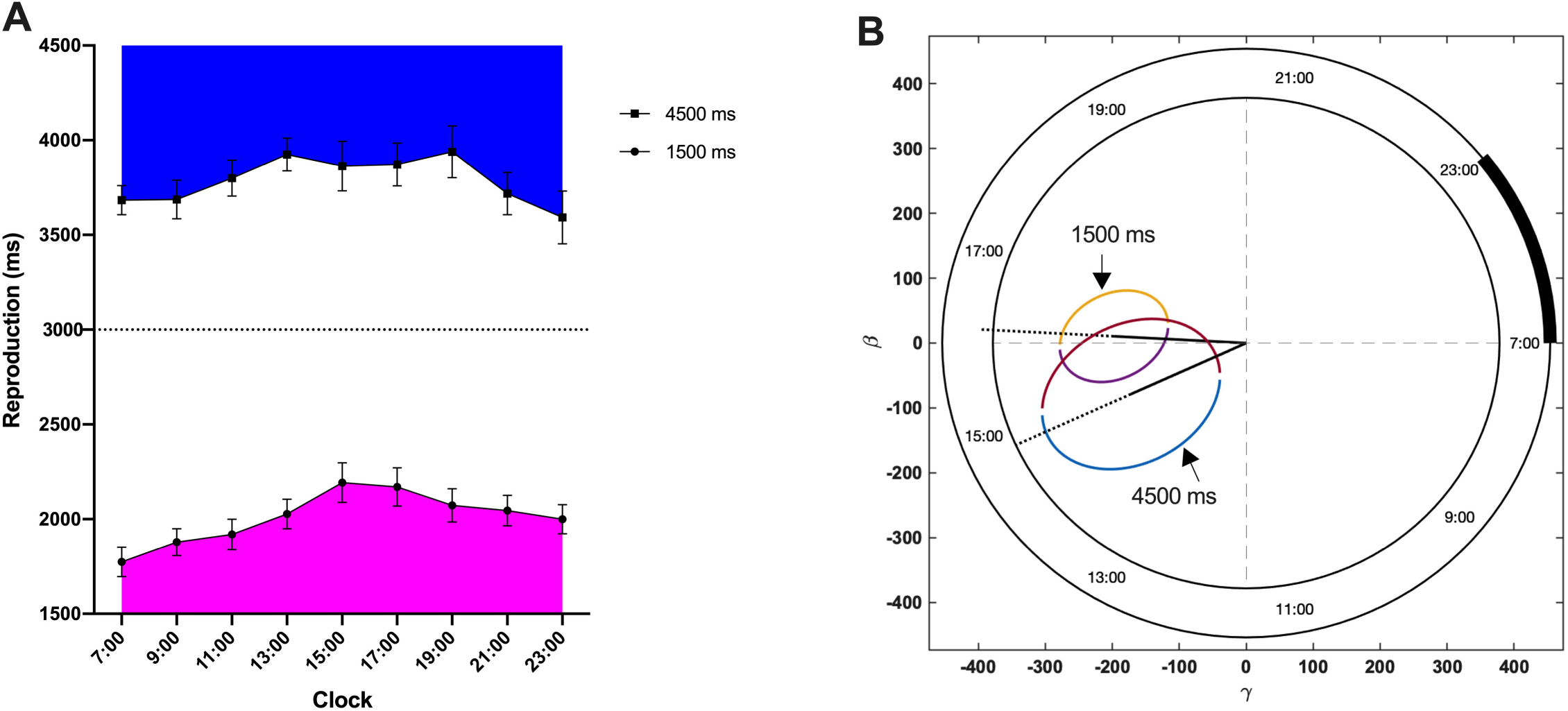
A. The diurnal rhythm of temporal reproduction (mean ± SEM). B. Polar coordinate of acrophases in two reproductions. β = Acosϕ; γ =-Asinϕ, A is the amplitude (a measure of half the extent of predictable variation within a cycle), ϕ is the acrophase (a measure of the time of overall high values recurring in each cycle). The regression model: Y= M+ A cos (ωt+ϕ), M is the MESOR (Midline Statistic Of Rhythm, a rhythm-adjusted mean). Corresponding values are: ‘lights on’, 7:00 (0°) and ‘lights off’, 23:00 (320°).

Abundant evidence from behavioral and neural studies show that 2-3 second window plays important roles in temporal processing. A typical conclusion has indicated that intervals that are longer than 2-3 s can no longer be perceived as a temporal-integration processing unit (Wittmann 1999). In this study, the reproductions have shown inversed estimations over and below 3 second window. Furthermore, the overestimated reproduction of 1500 ms showed smaller difference to the standard in the morning and evening and larger difference in the afternoon, and the underestimated reproduction of 4500 ms showed an inverse change (Figure 1.A). It suggested that two different processes exist for the duration-unit reproduction mechanism in the 2-3 second time window and vary with circadian rhythms. Temporal reproduction shows significant rhythms during the daytime, with acrophases of the performance occurring in the afternoon, but with the inversed mechanism of efficiency. No eccentricity effect in temporal production were observed, which indicated a homogenous temporal processing of reproduction shared in the perifoveal and peripheral visual field. This is consistent with the previous findings that the two attentional systems share the same time window(Bao, Pöppel et al. 2015).

Conclusively, the circadian rhythm of the temporal reproduction in a time window of 2 to 3 seconds plays an important role. The visual studies for the temporal processing not only consider the efficiency, such as eccentricity effect, but also should take circadian effect into account.

## Acknowledgements

This work was supported by XXX Foundation to Y.B. and the China Scholarship Council to the student W.W (CSC 201806020189) and X.Y (CSC 202108080312).

